# Fisetin extends lifespan in a murine model of recessive dystrophic epidermolysis bullosa

**DOI:** 10.1101/2024.11.23.625023

**Authors:** William C. Miller, Chloe L. Strege, Courtney M. Popp, Simone M. Intriago Tito, Cindy Eide, Christine L. Ebens, Megan Riddle, Chris Lees, Davis Seelig, Kacey Bui, Matthew J Yousefzadeh, John McGrath, Jakub Tolar

## Abstract

Recessive Dystrophic Epidermolysis Bullosa (RDEB) is a rare genodermatosis characterized clinically by extensive inflammation, cutaneous destruction, and fibrosis that demonstrates properties similar to rapid skin aging. As tissue ages, it accumulates cellular damage and exhaustion leading to a state of senescence. Cellular senescence is an aging or disease-related phenomenon of stable exit from the cell cycle that leads to an increased inflammatory phenotype. RDEB and other EB subsets of patients need an adjunct to or alternate therapy that addresses the issues of inflammation, pain, and pruritus. Fisetin is a safe, naturally occurring compound proven to be effective at sensitizing senescent cells to cell death and ameliorating senescence-associated inflammation. In this paper, we demonstrate fisetin’s ability to increase survival and reduce senescent cell burden in a hypomorphic mouse model of RDEB.

## Introduction

Recessive Dystrophic Epidermolysis Bullosa (RDEB) is a severe form of Epidermolysis Bullosa (EB), caused by mutations in the *Col7a1* gene [1]. The *Col7a1* gene is responsible for the coding of type VII collagen, a protein important for maintaining structural integrity in the basement membrane zone of the skin. Type VII collagen acts as an anchoring fibril between the epidermis and dermis [2]. The lack of functional type VII collagen results in mechanical fragility of the skin, causing debilitating chronic wounding, inflammation, pruritus, and scarring [3]. The devastating nature of RDEB, however, does not end here.

Significant fibrosis and inflammation around sites of chronic wounding predispose patients with RDEB to an increased risk of developing cutaneous squamous cell carcinoma (cSCC). Sadly, 90% of RDEB patients will develop cSCC by the age of 55, making it the major cause of death for these individuals [4]. In RDEB patients, the link between chronic wounding, with its associated inflammation, and cSCC development has significant supportive evidence [3]. A major driving mechanism of overactive inflammatory response is the phenomenon of cellular senescence.

Cellular senescence is a highly stable exit from the cell cycle and a typical stress response which can be caused by a vast array of factors. A senescent state can be induced by cellularly intrinsic or extrinsic factors such as radiation, chemotherapeutic drugs, oxidative stress, or genotoxic stress. Senescence is a dynamic progression that a cell undergoes starting with an initiation state that is controlled by the *p53/p21*^*Cip1*^ and *p16*^*Ink4a*^*/Rb* activation pathways, from which morphological and phenotypic changes cascade [5]. As cells enter long-term senescence, they begin to exhibit features such as increased cellular size, increased beta-galactosidase activity due to increased lysosomal content, and a specific secretory phenotype called the senescence-associated secretory phenotype (SASP) [6], which consists of a milieu of pro-inflammatory cytokines, interleukins, and growth factors that directly affect nearby cells [7].

The presence of senescent cells is not an inherently negative phenomenon, as this mechanism is needed transiently for organismal development, tissue remodeling, and cancer prevention [8-10]. However, prolonged late-stage senescence has been shown to be a major driver of aging and inflammation [11, 12]. Fibroblast senescence in skin tissue induced by aging was shown to directly impact keratinocyte differentiation and wound healing [13]. Interestingly, the skin of RDEB patients closely resembles the phenotypic and ultrastructural changes found in natural aging skin [14]. Data has also shown that senescence in the skin plays a major role in disrupting the epidermal-dermal junction, implicating it as a driving factor in the severity of the inflammatory phenotype of RDEB skin [15].

Reducing cell senescent burden has been shown to reduce aging symptoms such as inflammation and increase the proliferative capacity of stem and progenitor cells [16]. Compounds that selectively induce apoptosis of senescent cells are called senolytics [17]. Originally reported senolytics consisted of repurposed cancer chemotherapeutics: Navitoclax, Dasatinib, and Quercetin. Unfortunately, the use of these drugs can carry detrimental outcomes such as hemotoxicity as well as other generalized side effects linked with their use. In contrast, fisetin, a naturally occurring flavonoid, was discovered to have senotherapeutic properties and no known adverse side effects [18, 19]. Fisetin is a highly effective senolytic that induces apoptosis in senescent cells via inhibition of BCL-X_L_ [20]. Administration of fisetin to *Ercc1*^*-/Δ*^ mouse model of accelerated aging or naturally aged mice demonstrated significant reduction in the cell cycle inhibitors *p16*^*Ink4a*^ and *p21*^*Cip1*^ as well the proinflammatory molecules *Il-1β, Il-10, Il-6, Tnf-a, Cxcl2, Mcp1*, and *Pai1* in multiple tissues [18]. Because of its high senolytic capacity and significant reduction in inflammatory markers, fisetin was chosen to treat the fragile RDEB mouse model.

## Results

Fisetin administration to pregnant mothers and mutant pups was shown to increase the survival of RDEB pups. The demonstration of an increased cellular senescence burden in the RDEB pups warranted the usage of a compound that selectively clears senescent cells. Fisetin, due to its high senolytic capacity and lack of known negative side effects, was the chosen treatment. Fisetin’s impact on the RDEB phenotype was initially observed via longitudinal survivorship analysis. Given the decreased lifespan of RDEB pups, timed matings were established, and fisetin-dosed chow was administered to pregnant mothers at the time of separation from breeding males. Fisetin-treated chow (500 ppm) was available *ad libitum* to the treated group until the time of death. Litters were observed daily, and genotypes were confirmed at the time of natural death. The RDEB pups in litters treated with fisetin chow demonstrated a statistically significant increased survival compared to the untreated control group (p-value *≤*0.05) (Fig 1A). The fisetin group had a longer median survival of 8.5 days compared to the RDEB pups on the control diet with a median survival of 5 days (HR 2.21 95% CI 1.1 - 4.4). Alternatively, group characteristic survival demonstrated a 10-day successful survival rate of 48% in the treated group compared to 18% in the untreated group (Fig 1B).

**Figure 1.**
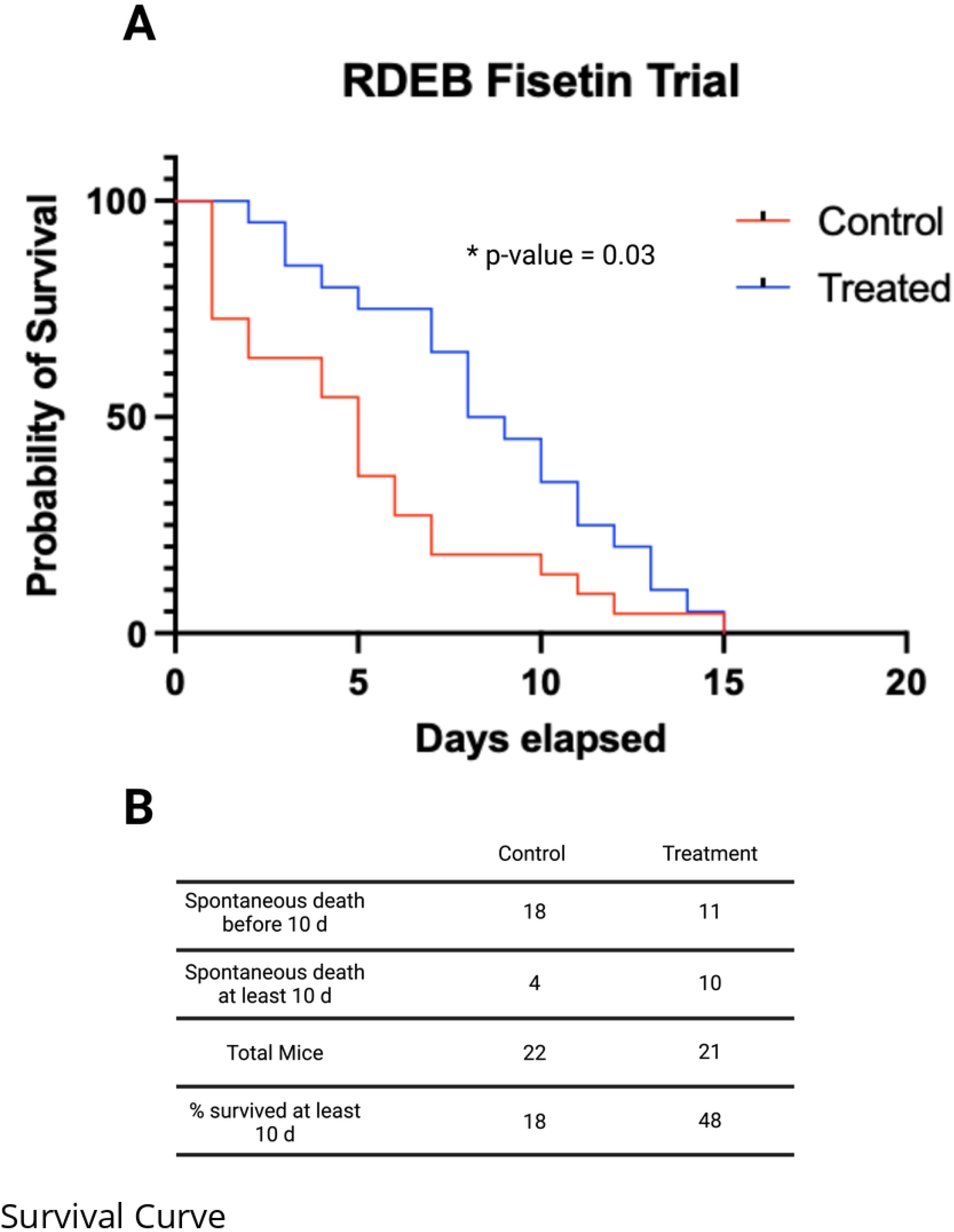
Fisetin increases the survival of RDEB pups. **A)** Kaplan-Meier survivorship curve of RDEB pups treated with fisetin compared to untreated RDEB pups. Fisetin-treated mice have a statistically significantly different probability of survival. Significance determined via Logrank (Mantel-Cox) survivorship analysis. *p-value ≤0*.*05* **B)** Distribution analysis of survivability to 10 days is statistically significantly different between treated and control groups. Significance determined via Chi-Squared Test of Independence. *p-value ≤ 0*.*05*

Fisetin administration reduces localized skin cellular senescent burden in RDEB pups. To assess fisetin’s impact on the levels of cellular senescence in RDEB, RT-qPCR analysis of skin tissue from treated and untreated RDEB pups was performed. Analysis of *p16*^*Ink4a*^ and *p21*^*Cip1*^ revealed a statistically significant reduction in *p16*^*Ink4a*^ in the fisetin-treated group (Fig 2). RT-qPCR of local skin SASP factor expression showed no statistically significant decrease in any of the measured transcripts. Systemic analysis of serum senescence and inflammatory markers showed no significant difference in *G-CSF, Eotaxin, GM-CSF, Il-2, Il-4, Il-3, Il-5, Il-6, Il-9, Il-10, Il-17, KC, MiP-1a*, or *RANTES*. Interestingly, *TNF-α* was statistically significantly higher in the serum of the fisetin treatment group compared to the untreated group. SA-βgal staining of treated and untreated RDEB skin showed foci of positive staining cells. However, there were no noticeable differences between the groups. *Col7a1* staining at 3, 13, and 21 days postpartum demonstrated no apparent increase in collagen VII expression between the fisetin-treated and untreated RDEB pups (Fig 3).

**Figure 2.**
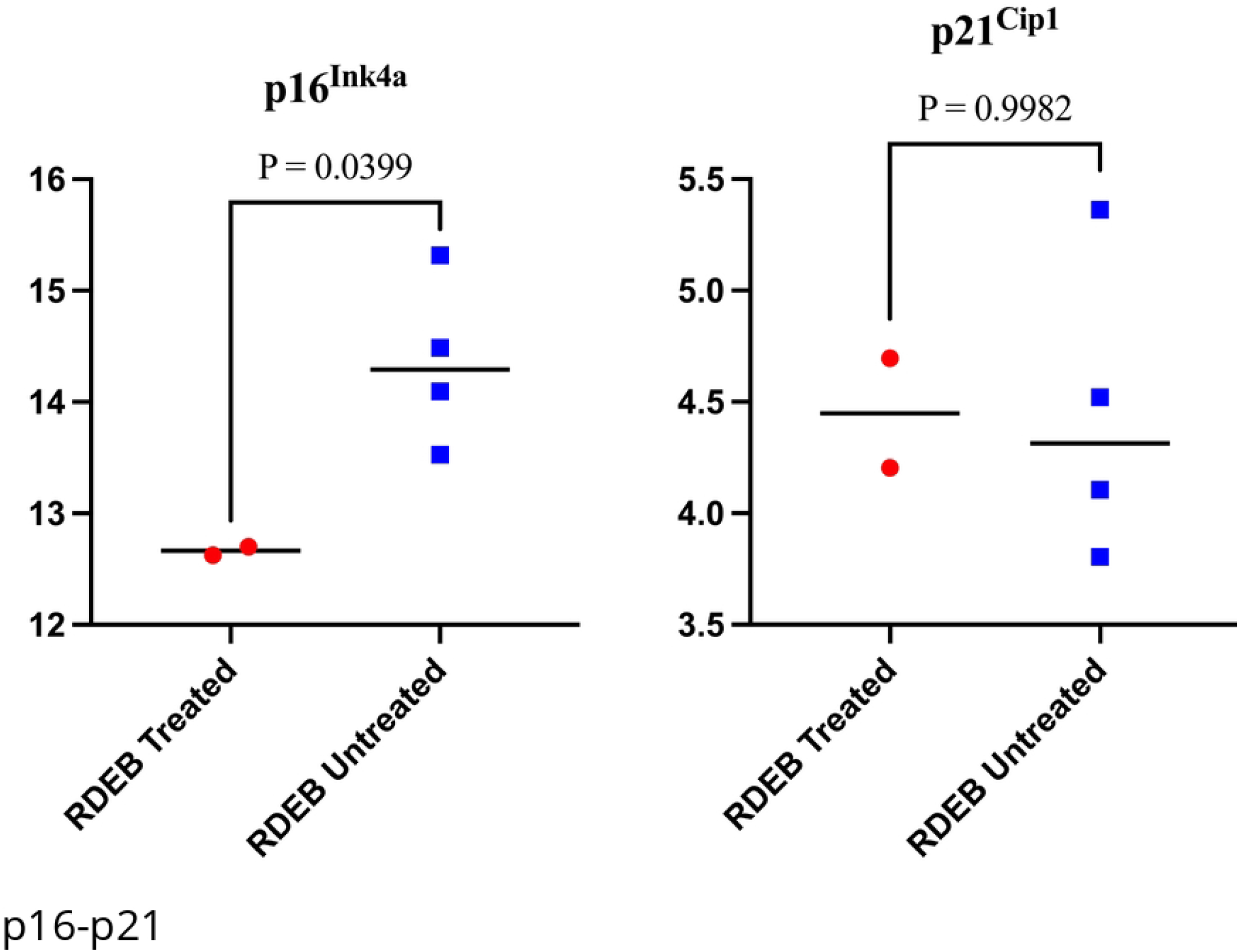
Fisetin reduces local skin expression of p16^Ink4a^. RT-qPCR analysis of bulk skin tissue comparing *p16*^*Ink4a*^ and *p21*^*Cip1*^ expression in RDEB treated and untreated 3-day old pups. Significance determined via two-tailed student’s t-test. *\* p ≤ 0*.*05*

**Figure 3.**
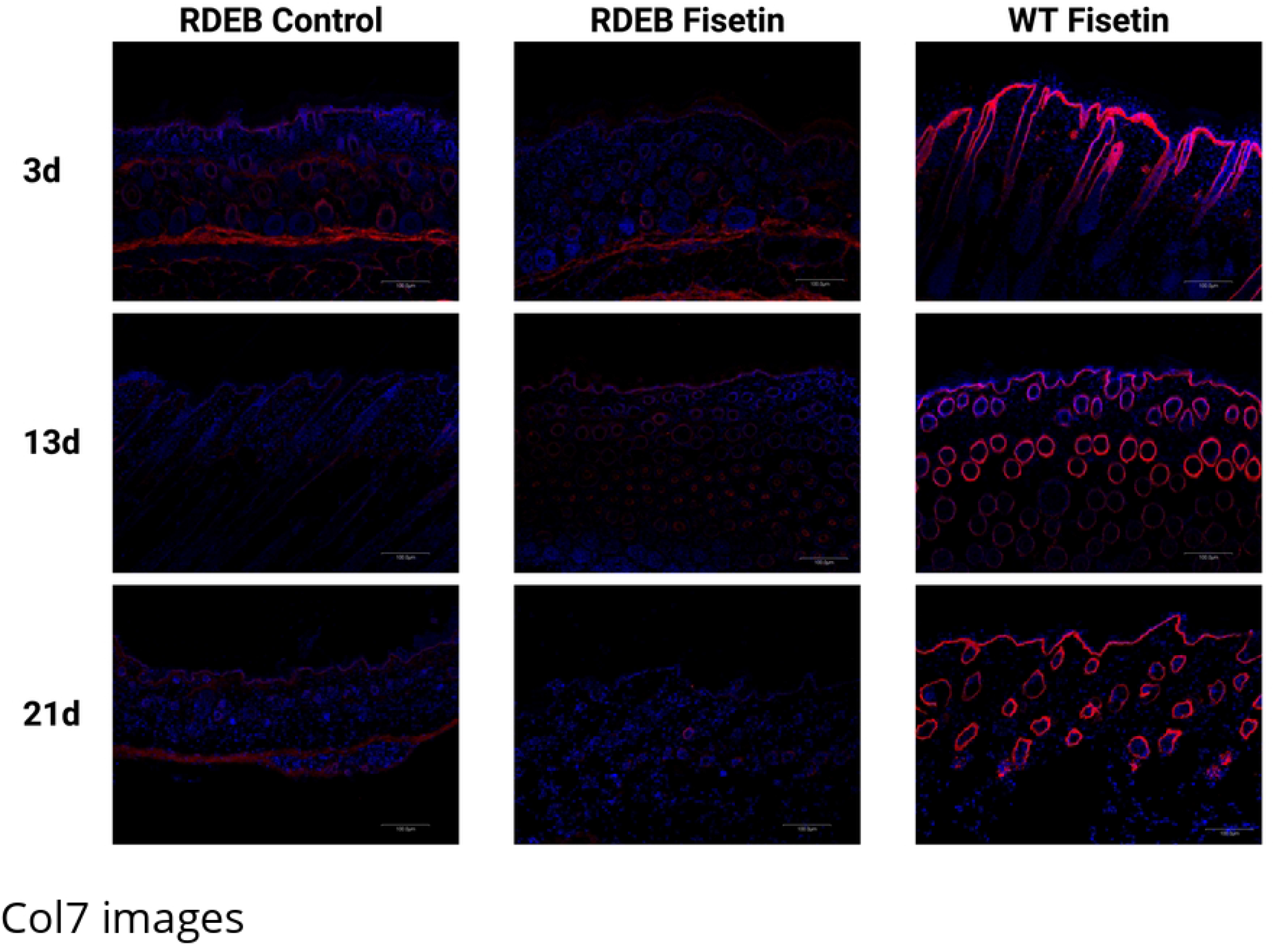
Immunofluorescence demonstrates no increase of collagen VII staining between treated and untreated groups. Immunofluorescence of collagen VII across all groups at 3, 13, and 21 days captured at 20x magnification.

## Discussion

RDEB is a destructive inherited skin disorder with resultant severe mechanical fragility of the skin, systemic inflammation, and an inevitable progression to lethal cSCC. There remains a paucity of safe and effective treatments targeting inflammation and associated symptoms for this disease. Literature continues to emerge characterizing the phenomenon of cellular senescence as a driver of chronic inflammation associated with aging and multiple disease states. In this study, we demonstrated a senescent phenotype exacerbated by RDEB and the positive benefits of its targeted reduction.

In a cumulative analysis of RDEB senescence, the cell cycle inhibitor *p16*^*Ink4a*^ displayed the largest difference when compared to WT mice. *p16*^*Ink4a*^ is associated with DNA damage due to telomere dysfunction and is a specific marker of cellular senescence. Alternatively, *p21*^*Cip1*^ is known to be elevated during senescence, but it may be transiently elevated for numerous non-senescent related reasons [21]. The loss of *p16*^*Ink4a*^ results in reduced senescence burden [22].

Additionally, mutations in *CDKN2A*, for which *p16*^*Ink4a*^ is one of the gene products, are among the leading causes of RDEB-cSCC [23]. Given the potential role of *p16*^*Ink4a*^ in RDEB symptom severity and cSCC development, it is an ideal therapeutic target for senolytic therapy. Our experiments demonstrated that *p16*^*Ink4a*^ was significantly reduced with the treatment of fisetin.

While the local expression increase of SASP factors in RDEB skin was insignificant, the same factors were found to be systematically increased, demonstrating significant organismal inflammatory involvement. The overwhelming nature of inflammation in a disease such as RDEB represents a significant problem, but the ability of a single therapy to significantly reduce molecular markers of inflammation may be unrealistic. However, any reduction of inflammation may demonstrate impactful clinical benefits on the organismal level. This is supported by the significant increase in survival exhibited by the fisetin-treated RDEB pups.

Fisetin provides a safe, affordable, and effective treatment to increase lifespan and reduce symptom severity in RDEB mice. Because the exact involvement of cellular senescence in chronic wounding is not completely understood, further spatial and temporal analysis of the skin microenvironment during persistent wounding and subsequent healing is required for optimal therapy. While the molecular mechanism of fisetin in RDEB is unclear, the positive impact fisetin has demonstrated in this hypomorphic mouse model warrants further investigation into its therapeutic potential. The logical next step in experimentation, is to examine the effect fisetin has in the prevention or reduction in cSCC development and institute long-term studies on the usage of senolytics or senescent cell ablation to improve the phenotype of RDEB mice.

## Methods

### Murine Model

Mouse research was approved by the Institutional Animal Care and Use Committee (IACUC) at the University of Minnesota (UMN) under protocol number 2106-39156A. All mice were housed in standard, small animal research conditions per University IACUC guidelines with a 12-hour light:dark cycle, unlimited access to food and water, temperatures ranging from 22-24°C, and humidity levels ranging from 30% to 32%. The cages were housed on the rack furthest from the door and laminar flow hood, with all cages on the side of the rack facing the corner wall. All animals were housed in static Allentown Mouse 75Jag Cages (Allentown, Allentown, NJ) with ALPHA-Dri PLUS bedding and Enviro-dri nesting material (Shepherd Specialty Papers, Amherst, MA). Enrichment was also provided to all cages in the form of steam-sterilized cardboard glove boxes, cut in half. To decrease outside stressors and potential discomfort, the handling of mice and their cages was limited to 2 laboratory staff members trained in gentle handling techniques. These staff members were responsible for cage changes and twice daily monitoring of the animals’ health, behavior, and survival status. If health or behavioral concerns were to arise that involved medical intervention, the laboratory members would partner with veterinary staff in the handling and treatment of animals. If the following humane endpoints were observed, it would result in immediate euthanasia: 20% weight loss, inability to reach food or water, not nursing in the case of pre-weaned mice, and failure to grow. While these humane endpoints are designed to prevent animals from reaching a moribund state, if an animal was found moribund, defined by UMN IACUC as unresponsive to gentle physical stimulation, the animal would also be immediately euthanized.

A previously established type VII collagen hypomorphic mouse model kept on a C57BL/6:129sv background was used (Fritsch, 2008). Timed matings were established and at the time of separation, approximately 24-72 hours later, females were given their respective diets. The treatment group was given unlimited access to Teklad 2020 chow (Envigo, Madison, WI) prepared with 500 ppm (500 mg/kg) of Fisetin (Indofine Chemical Co., Hissborough, NJ) by Envigo Co. The control group was given unlimited access to Teklad 2919 irradiated chow (Envigo, Madison, WI). As previously stated, litters were checked twice daily to establish survival data with genotypes confirmed at the time of death via PCR using a previously established protocol [24]. A total of 43 diseased type VII collagen hypomorphic mouse pups were observed for the duration of this experiment. Of these 43 animals, none met the humane endpoints listed above, and therefore did not require immediate euthanasia. All were found dead at varying times between birth and day 15 post-birth. All rodents sacrificed for tissue collection were first euthanized using the UMN IACUC approved procedure of decapitation. For tissue harvest, pups were taken at 3, 13, and 21 days and genotypes were confirmed via PCR.

### Senescence-Associated β-Galactosidase Staining

Senescence-associated β-galactosidase activity in skin tissue was performed using a standard SA-βgal staining procedure [25]. Skin tissue samples were fixed in 10% neutral buffered formalin for 4 hours, placed in 30% sucrose for 24 hours, and embedded in O.C.T prior to cryosectioning at 6 microns. Tissue sections were stained with an X-gal containing solution, where the pH was adjusted with NaOH and HCl as necessary to reach a pH of 4.5 [26]. Samples were then imaged using bright field microscopy on an EVOS XL Core Imaging System at 4X, 10X, 20X and 40X magnification.

### Immunofluorescence Staining

RDEB untreated, RDEB treated, and wildtype treated murine skin tissue samples were frozen in optimal cutting temperature (OCT, Sakura Finetek USA, Torrance,CA). The blocks were sectioned at 6 microns on a cryostat. Slides were fixed in room temperature acetone for 5 minutes. Tissue sections were blocked with 10% Normal Donkey Serum for 1 hour (Jackson Immunoresearch, West Grove, PA). Primary antibody collagen VII (1:200 LifSpan BioSciences, Seattle, WA) was incubated overnight at 4°C. Slides were washed with 1x PBS. Secondary donkey anti-rabbit Cy3 (Jackson Immunoresearch, West Grove, PA 1:500) was applied for 1 hour at room temperature. Slides were washed with 1xPBS and then cover slipped with hard set DAPI, 4,6-diamidino-2-phenylindole, (Vector Labs, Burlingame, CA). Slides were examined by confocal fluorescence microscopy (Olympus BX61, Olympus Optical, Tokyo,Japan).

### Tissue Histopathology

Skin tissue was harvested from euthanized mice and fixed in 10% neutral buffered formalin for 24 hours, then transferred to 70% EtOH. Tissues were then sent to the Comparative Pathology Shared Resource (CPSr) at the University of Minnesota where they were paraffin-embedded, sectioned, and mounted onto slides under the direction of Dr. Seelig for H&E staining and veterinary pathology diagnosis.

### RT-qPCR

Skin tissue was harvested from euthanized mice, snap-frozen on dry ice, and stored at - 80°C until use. RNA extraction from tissue was performed using the Qiagen RNEasy mini kit (Qiagen Cat. # 74104), reverse transcription (RT) was performed using Thermo Fisher Superscript VILO (Catalog: 11754050), and RT-qPCR was performed using Thermo Fisher TaqMan Gene Expression Master Mix (Catalog: 4369016). TaqMan probes were used for RT-qPCR targeting *p16*^*Ink4a*^ (Mm00494449_m1), *p21*^*Cip1*^ (Mm04205640_g1), *IL-1β* (Mm00434228_m1), *IL-6* (Mm00446190_m1), *IL-10* (Mm01288386_m1), *CCL2* (Mm00441242_m1), *TNF-α* (Mm00443258_m1) and *GAPDH* (Mm99999915_g1). For RT-qPCR, 20uL reactions were prepared using 1uL of 20X from each probe and 10 ng of cDNA. Each RT-qPCR was run in triplicate and analyzed using the ΔΔCT method, averaging the values across triplicates. *Gapdh* was used as an internal control.

### Serum Analysis

At the time of euthanasia, blood was collected in a microcentrifuge tube and allowed to sit at room temperature until a clot was formed. Samples were then centrifuged for 20 minutes at 2000 x g and the supernatant was removed. Serum samples were then submitted to the Cytokine Reference Laboratory at the University of Minnesota for analysis (CRL, University of Minnesota). This is a CLIA’88 licensed facility (license #24D0931212) under the direction of the American Board of Medical Laboratory Immunology Certified Dr. Mortari. Commercially available kits and reagents were used. Samples were analyzed for mouse-specific cytokines and chemokines using the Luminex platform and done as a multi-plex. The magnetic bead set (cat. #MCYTOMAG-70K) was purchased from EMD Millipore, St. Charles MO. Samples were assayed according to manufacturer’s instructions. The beads were read on a Luminex instrument (MAGPIX). Samples were run in duplicate and values were interpolated from 5-parameter fitted standard curves.

### Statistics

For survivorship analysis, survival data from group mice was taken and analyzed for outliers using the ROUT method with Q = 1%. Cleaned data was analyzed using the logrank (Mantel-Cox) method with no censored events. RT-qPCR and serum cytokine/chemokine analysis was performed using a two-tailed student’s t-test with a significance threshold of 5% (0.05). 10-day relative survival percentages compared using Chi-Squared Test of Independence. All statistical analyses were performed in GraphPad Prism v 9.4.1 (458).

## Abbreviations

RDEB: recessive dystrophic epidermolysis bullosa
EB: epidermolysis bullosa
cSCC: cutaneous squamous cell carcinoma
SASP: senescence-associated secretory phenotype
IACUC: Institutional Animal Care and Use Committee

## Author Contributions

The authors confirm contribution to the paper as follows: study conception and design: William C. Miller; data collection: William C. Miller, Chloe L. Strege, Simone M. Intriago Tito, Courtney M. Popp, Megan Riddle, Davis Seelig; analysis and interpretation of results: William C. Miller, Chloe L. Strege, Megan Riddle, Chris Lees, Davis Seelig, Matthew J Yousefzadeh; draft manuscript preparation: William C. Miller, Chloe L. Strege, Courtney M. Popp, Cindy Eide, Christine L. Ebens, Megan Riddle, Kacey Bui, Matthew J Yousefzadeh, John McGrath, Jakub Tolar. All authors reviewed the results and approved the final version of the manuscript.

## Acknowledgments

We would like to acknowledge discussions with members of the University of Minnesota Institute on the Biology of Aging and Metabolism (iBAM) and the Department of Biochemistry, Molecular Biology, and Biophysics.

